# Ipsilateral finger representations are engaged in active movement, but not sensory processing

**DOI:** 10.1101/285809

**Authors:** Eva Berlot, George Prichard, Jill O’Reilly, Naveed Ejaz, Jörn Diedrichsen

## Abstract

Hand and finger movements are mostly controlled through crossed corticospinal projections from the contralateral hemisphere. During unimanual movements, activity in the contralateral hemisphere is increased while the ipsilateral hemisphere is suppressed below resting baseline. Despite this suppression, unimanual movements can be decoded from ipsilateral activity alone. This indicates that ipsilateral activity patterns represent parameters of ongoing movement, but the origin and functional relevance of these representations is unclear. Here, we asked whether human ipsilateral representations are caused by active movement, or whether they are driven by sensory input. Participants alternated between performing single finger presses and having fingers passively stimulated, while we recorded brain activity using high-field (7T) functional imaging. We contrasted active and passive finger representations in sensorimotor areas of ipsilateral and contralateral hemispheres. Finger representations in the contralateral hemisphere were equally strong under passive and active conditions, highlighting the importance of sensory information in feedback control. In contrast, ipsilateral finger representations were stronger during active presses. Furthermore, the spatial distribution of finger representations differed between hemispheres: the contralateral hemisphere showed the strongest finger representations in Brodmann area 3a and 3b, while the ipsilateral hemisphere exhibited stronger representations in premotor and parietal areas. This suggests that finger representations in the two hemispheres have different origins – contralateral representations are driven by both active movement and sensory stimulation, whereas ipsilateral representations are mainly engaged during active movement. This suggests that a possible contribution of the ipsilateral hemisphere lies in movement planning, rather than in the dexterous feedback control of the movement.

**Significance statement:** Movements of the human body are mostly controlled by contralateral cortical regions. However, activity in ipsilateral sensorimotor regions is also modulated during active movements. The origin and functional relevance of these ipsilateral representations is unclear. Here we used high-field neuroimaging to investigate how human contralateral and ipsilateral hemispheres represent active finger presses and passive finger stimulation. We report that while the contralateral hemisphere was equally strongly recruited during active and passive conditions, the ipsilateral hemisphere was mostly recruited during active movement. We propose that the ipsilateral hemisphere may play a role in bilateral movement planning.

## Introduction

The primate hand is controlled mainly by descending projections from the motor areas in the contralateral cerebral hemisphere (Brinkman and Kuypers, 1973). While the hand also receives input from ipsilateral motor regions through uncrossed corticospinal projections, these projections lack the capacity to produce overt movement (Soteropoulos et al., 2011). If, and to what degree the ipsilateral hemisphere directly or indirectly contributes to hand movements is currently debated (Chen et al., 1997; Verstynen et al., 2005). It is clear, however, that neural activity in ipsilateral motor regions is modulated during hand movements. Overall, there is a global suppression of activity as evidenced by a reduction in BOLD (blood-oxygen-level-dependent) signal measured using functional magnetic resonance imaging (fMRI) (Verstynen et al., 2005; Cramer et al., 2011). Below this suppressive effect, there are clear task-specific changes. For example, one can decode the identity of the moved effector (e.g. finger) from ipsilateral activity alone (ECoG: Scherer et al., 2009; Fujiwara et al., 2017; fMRI: Diedrichsen et al., 2013). These ipsilateral activity patterns appear to be weaker, but otherwise identical versions of the pattern elicited by movement of the mirror-symmetric finger in the opposing hand (Diedrichsen et al., 2013, 2017). Altogether, these studies show that the ipsilateral hemisphere *represents* aspects of finger movements. The origin and functional relevance of these representations, however, remain unclear.

One puzzle regarding the function of these ipsilateral representations is whether they reflect processes involved in active motor planning and execution, or whether they are a consequence of re-afferent sensory input. In the contralateral hemisphere, passive somatosensory stimulation of individual fingers has been shown to evoke activity patterns that are very similar to those associated with active finger movements (Wiestler et al., 2011). This is even the case on the single-finger level; cortical patches that are especially activated by movement of the index finger are also activated by index finger stimulation. The tight match between tuning for active and passive conditions is unsurprising, given how important accurate sensory information from the fingertip is for fine control of each digit (Augurelle et al., 2003; Pruszynski et al., 2016). Overall, this architecture indicates that the main role of motor cortex is *feedback* control (Scott, 2004).

Here we ask whether ipsilateral sensorimotor cortex plays a role in the fine feedback control of finger movements. If so, we should see that ipsilateral representations can also be activated by passive sensory stimulation. Indeed, we would expect that passive finger stimulation recruits ipsilateral finger-specific circuits to approximately the same degree as active finger presses, as they do in the contralateral sensorimotor cortex. Alternatively, if the ipsilateral hemisphere is primarily recruited during movement planning, we would predict that ipsilateral representations are more pronounced during active presses, and either weaker or absent during passive finger stimulation.

To test between these two possibilities, we used high-field fMRI (7T) to measure ipsilateral activity patterns during active single finger presses and passive finger stimulation. We contrasted the overall activity during active and passive conditions in both the contralateral and ipsilateral hemisphere. Using multivariate pattern analysis, we also analyzed how strongly different conditions activated finger-specific circuits, i.e. the degree to which finger information is represented in these areas (Diedrichsen and Kriegeskorte, 2017). This analysis allowed us to determine the extent to which representations in the contralateral and ipsilateral motor areas are driven by sensory input alone (passive condition), or by a combination of sensory input and active planning and execution processes (active condition). We further examined these representations using a fine-grained analysis across the subfields of the sensorimotor cortices.

Overall, we found that active and passive conditions recruited contralateral and ipsilateral sensorimotor areas differently. While contralateral finger representations were equally strong for active and passive conditions, the corresponding ipsilateral finger representations around the central sulcus were stronger for the active than passive condition. This differential recruitment of contralateral and ipsilateral motor areas suggests an involvement of ipsilateral areas in active motor planning.

## Method

### Participants

Seven volunteers participated in the experiment. The average age was 26.1 years (SD = 2.5 years), and the sample included 4 women and 3 men. All participants were right-handed. The experimental procedures were approved by the Ethics Committee of University College London and Oxford University.

### Apparatus

Participants placed their two hands on an MRI-compatible keyboard (Fig. 1A), which was positioned on their lap, secured with a foam pillow. The keyboard had 10 elongated keys, with a groove for each fingertip. Force applied during finger press execution was measured with force transducers mounted underneath each key. The keys were non-movable and therefore finger presses were not associated with overt movements. Nonetheless, these isometric presses still involved voluntary activation of muscles, as well as sensory feedback from the pressure on the fingertip. To generate a sensory stimulation protocol that was matched as closely as possible to the sensory input during active finger presses, we applied isometric force presses through pneumatic pistons embedded in each key of the keyboard. Upward movement of the finger was prevented by a stiff foam pad which held the fingers securely in place (Fig. 1B). The force to the fingertip in the passive condition was closely matched to that in the active condition by generating force pulses at the same inter-press interval and with the same average peak force as those produced during the active condition. The mean peak force was 4.3N in the active and 4.5N in the passive condition. Therefore, the two conditions differed mainly in terms of the motor command (i.e. the efference), and were matched as closely as possible in terms of sensory afference. It is of course never possible to exactly match sensory feedback across active and passive conditions, as the efferent outflow itself will alter the incoming sensory information (Blakemore et al., 1999). Therefore, our conclusions on the source of representation did not rely on a direct comparison of passive and active conditions, but rather on a difference in their relative weighting in contra-vs. ipsilateral sensorimotor regions.

**Fig. 1.**
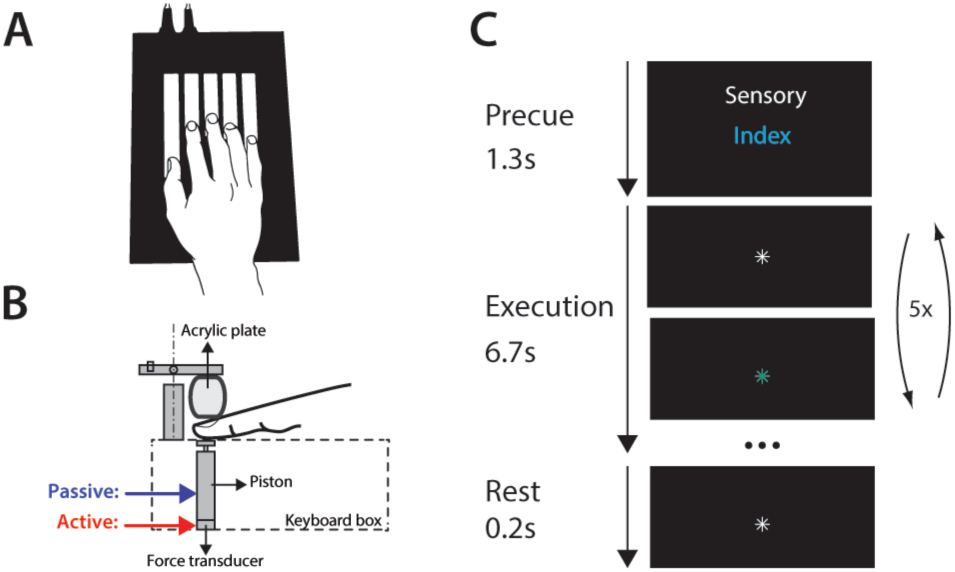
Apparatus and experimental design. (**A**) Keyboard used in the task – the left hand was positioned on a mirror-symmetric keyboard. (**B**) An adjustable foam pillow was sitting on the top of each finger, preventing any overt finger motion. In the active condition, participants pressed one of the keys and the force applied was recorded through the force transducer. In the passive condition, the force was applied to the finger using a pneumatic piston. (**C**) Each trial started with a cue denoting which condition and finger are implicated in the trial. This was followed by a warning press to the finger, after which each participant either received five finger presses (in the passive condition), or pressed the key five times. Each trial lasted for a total of 8.2 seconds. Both active and passive conditions involved only the right hand.

### Experimental design

We employed a slow event-related design, randomly intermixing active and passive conditions in each imaging run. Every trial lasted for 8.2 seconds, during which participants either performed five isometric presses with one of the fingers (active condition) or had a finger stimulated five times (passive condition). Both conditions involved only the right hand. Each trial was divided into the instruction phase (1.3 s) and the execution phase (6.7 s). First, the instructional cue was presented on the screen, specifying which finger is to be pressed or stimulated (e.g. Sensory / Index, Fig. 1C). Additionally, a warning press was applied to the finger which was to be pressed or stimulated. Afterwards, participants performed five presses, or had force applied to their finger five times, while fixating on a central cross. For every press with the correct finger, the central fixation point turned green, whereas it turned red for an incorrect press. The central fixation then turned white again to trigger the next press. Each run contained three repetitions of each of ten conditions (five fingers in passive / active tasks), and there were seven or eight imaging runs per participant. Five rest phases of 13 to 16 seconds each were randomly interspersed in each imaging run to obtain a reliable estimate of baseline activation.

### Image acquisition

Data was acquired on a 7 Tesla Siemens Magnetom scanner with a 32-channel head coil. An anatomical T1-weighted scan was acquired using a magnetization-prepared rapid gradient echo sequence (MPRAGE) with voxel size of 0.7 mm isotropic (field of view = 224×224×180 mm). Functional data was acquired in 7-8 runs (depending on the participant), using a 2D echo-planar imaging sequence (GRAPPA 2, repetition time [TR] = 3.0 s, echo time [TE] = 25 ms). We acquired 47 slices with isotropic voxel size of 1.4 mm.

### First-level analysis

Functional data were analysed using SPM12 and custom-written Matlab code. Differences in acquisition timing of slices were corrected for by aligning all slices to the middle slice of each volume. Functional images were corrected for geometric distortions using fieldmap data, and aligned to the first image of the first run, resulting in correction for head movements during the scan (3 translations: x, y, z directions and 3 rotations: pitch, roll, and yaw). Finally, the data was co-registered to the anatomical scan. No smoothing or normalisation to an atlas template was performed at this stage.

Preprocessed data were analysed using a general linear model. For each trial type, we defined one regressor per imaging run, resulting in 10 regressors per run (5 fingers in passive / active conditions). The regressor was a boxcar function which started with the beginning of the trial and lasted for the trial duration. This function was convolved with a hemodynamic response function, with a time-to-peak of 4.5s, manually adjusted to best fit the average timeseries. The analysis resulted in one activation estimate (beta image) for each of the ten conditions per run. We calculated average percent signal change for the passive and active conditions (averaged across all fingers) as the mean evoked response relative to the baseline in each run, averaged across runs.

### Surface-based analysis and searchlight approach

To carefully characterize activation patterns across different cortical areas, we obtained a reconstruction of individual subjects’ cortical surfaces using Freesurfer (Dale et al., 1999). All individual surfaces were aligned to the symmetrized atlas template of Freesurfer (using xhemireg, Fischl et al., 1999) via spherical registration.

To detect finger-specific representations for the active and passive conditions across the cortex (see multivariate analysis), we used a surface-based searchlight approach (Oosterhof et al., 2011). For each surface node, we selected a surrounding circular region of 120 voxels (i.e. in 3D volume), which on average resulted in a searchlight radius of 6.5 mm. To avoid contamination of signals across the central sulcus, we excluded all voxels that contained gray matter from the other side. We extracted the activation estimates (betas) of selected voxels from the first-level analysis and then computed the dissimilarity between activity patters for the passive and active finger pairs (see below). The resulting distance was assigned to the center of the searchlight sphere. By moving the searchlight across the cortical surface, we obtained a map of distances for active and passive condition patterns, representing how well each patch of cortex represented individual finger active and passive conditions.

### Regions of interest (ROI) and cross-section

To compare finger representations across different subfields of the sensorimotor cortex, we defined seven regions of interest (ROIs). The ROIs were defined using anatomical maps derived post-mortem histology that were aligned to the cortical surface atlas (Fischl et al., 2008). Each cortical node was assigned to the region that had (across analyzed brains) the highest probability. Primary motor cortex (M1), or Brodmann area 4, was split into anterior (BA4a) and posterior (BA4p) components. To exclude mouth and leg representations, we included only cortical nodes within a 2.5 cm distance from the hand knob. ROIs for primary somatosensory cortex (S1) were Brodmann areas 3a, 3b, 1, and 2. Additionally, the premotor cortex was defined as the lateral aspect of Brodmann area 6 (BA6).

We performed the analysis on percent signal change and distance estimates (see multivariate analysis) for cortical surface patches in a cross-section across the surface sheet, running from the rostral end of BA6 to the posterior end of BA2. For the pattern component modelling analysis (described below), we used all voxels within each ROI, and further joined BA4a and BA4p into BA4, and BA3a and BA3b into BA3.

### Multivariate analysis

The overall activation across fingers does not provide insight into finger-specific processes (i.e. finger representations; Diedrichsen et al., 2013). While finger representations can be visualized in terms of their rough somatotopic arrangement on the cortical surface (Indovina and Sanes, 2001; Wiestler et al., 2011), a fuller description can be obtained by taking into account the entire fine-grained activity pattern for each finger (Ejaz et al., 2015). We therefore calculated distances between activation patterns for different fingers, separately for each subject. We first standardized the beta-image for each voxel by dividing it by the standard deviation of that voxel’s residual, obtained from the first-level GLM. Such univariate prewhitening has been shown to increase the reliability of distance estimates as compared to non-standardized images (Walther et al., 2016). For active and passive conditions separately, we then calculated the crossvalidated squared Mahalanobis distance (crossnobis estimator, Nili et al., 2014; Walther et al., 2016; Diedrichsen and Kriegeskorte, 2017) between each finger pair. Because the expected value of this estimator is zero if the two conditions only differ by measurement noise, the crossnobis estimate can be used to test whether an area “*represents*” a certain parameter by testing it against zero (Diedrichsen et al., 2016).

### Pattern component analysis

To quantify the correspondence between active and passive activity patterns, we used pattern component modelling (PCM; Diedrichsen et al., 2017). A naïve way to assess the correlation would be to simply correlate corresponding finger patterns, after subtracting the mean pattern, for the passive and active condition. However, the raw correlations severely underestimate the true correlation between patterns as the correlations are lowered by measurement noise. Even cross-validated correlations are severely biased (see example 2, Diedrichsen et al., 2017). Instead, we can use PCM to test between different models on the strength of the correlation between the finger-specific patterns in the active and passive condition: a *‘null’ model* where active and passive conditions are unrelated, a *‘flexible correlation’ model* where the two conditions share some correlation, and a *‘perfect correlation’ model* in which the passive finger-specific patterns are simply scaled version of the active patterns. We compared these models by calculating for each subject the log-Bayes factor of the flexible and perfect model against the null model. Subsequent group inferences were performed using parametric statistics (t-test) on the individual log-Bayes factors.

### Statistical analyses

To statistically assess how activity or distances differ between conditions in either hemisphere, we performed a condition x ROI ANOVA, followed by post-hoc t-tests on distance estimates of passive and active conditions in each region individually. To directly contrast the distance estimates of the two conditions across the two hemispheres, we conducted a hemisphere x condition ANOVA. We further quantified the spatial distribution of distances across regions of the two hemispheres using a hemisphere x ROI ANOVA on estimates of distances in the active condition. To statistically assess the correspondence between active and passive patterns, we contrasted the obtained correlation estimates against 0 using one-sample t-tests, and conducted a model type x ROI ANOVA on log-Bayes factors of the flexible and perfect correlation models. Our ANOVAs were followed by post-hoc t-tests, using Bonferroni correction for multiple comparisons for adjusting the significance value. Given the small sample size (N=7), we replicated each test using non-parametric statistics (rank-sum test, not reported here), which yielded qualitatively similar results.

## Results

### Contralateral finger representations are equally strong in active and passive conditions

Before looking at the contribution of sensory and motor processes to the ipsilateral representations, we carefully quantified the passive and active finger representations in the contralateral hemisphere. As a first proxy for contralateral recruitment during the two conditions, we investigated the overall BOLD activation across sensorimotor regions. The sensory input was similar in both tasks, but the active condition additionally required planning and initiation of the press. These additional motor demands were predicted to evoke higher levels of activation in the active compared to the passive task. Figure 2A shows the percent signal change on the flattened cortical surface related to the active (red) and passive condition (blue). Both conditions evoke activity in highly overlapping cortical patches (purple). For statistical evaluation, we used a series of anatomically defined ROIs, running from premotor cortex (BA6) posterior into BA2 (separated by dashed white lines), and tested the evoked activity of each region against zero with a one-sample t-test. Significance at *p*<0.001 was reached in all subfields for both passive and active conditions (blue and red bars in Fig 2C). To examine differences between active and passive conditions, we performed a condition x ROI ANOVA. Both the main effects of condition and ROI were significant (condition: *F*_(1,6)_=23.791, *p*=0.0028, ROI: *F*_(1,6)_=4.833, *p*=9.0e^-4^), as was the interaction between them (*F*_(1,6)_=8.19, *p*=1.3e^-5^). Post-hoc t-tests comparing activation during passive and active conditions within each ROI revealed that the active condition elicited higher activation than the passive one in every region (Bonferroni-corrected significance level: *p*=0.0071 – black stars in Fig. 2C).

**Fig. 2.**
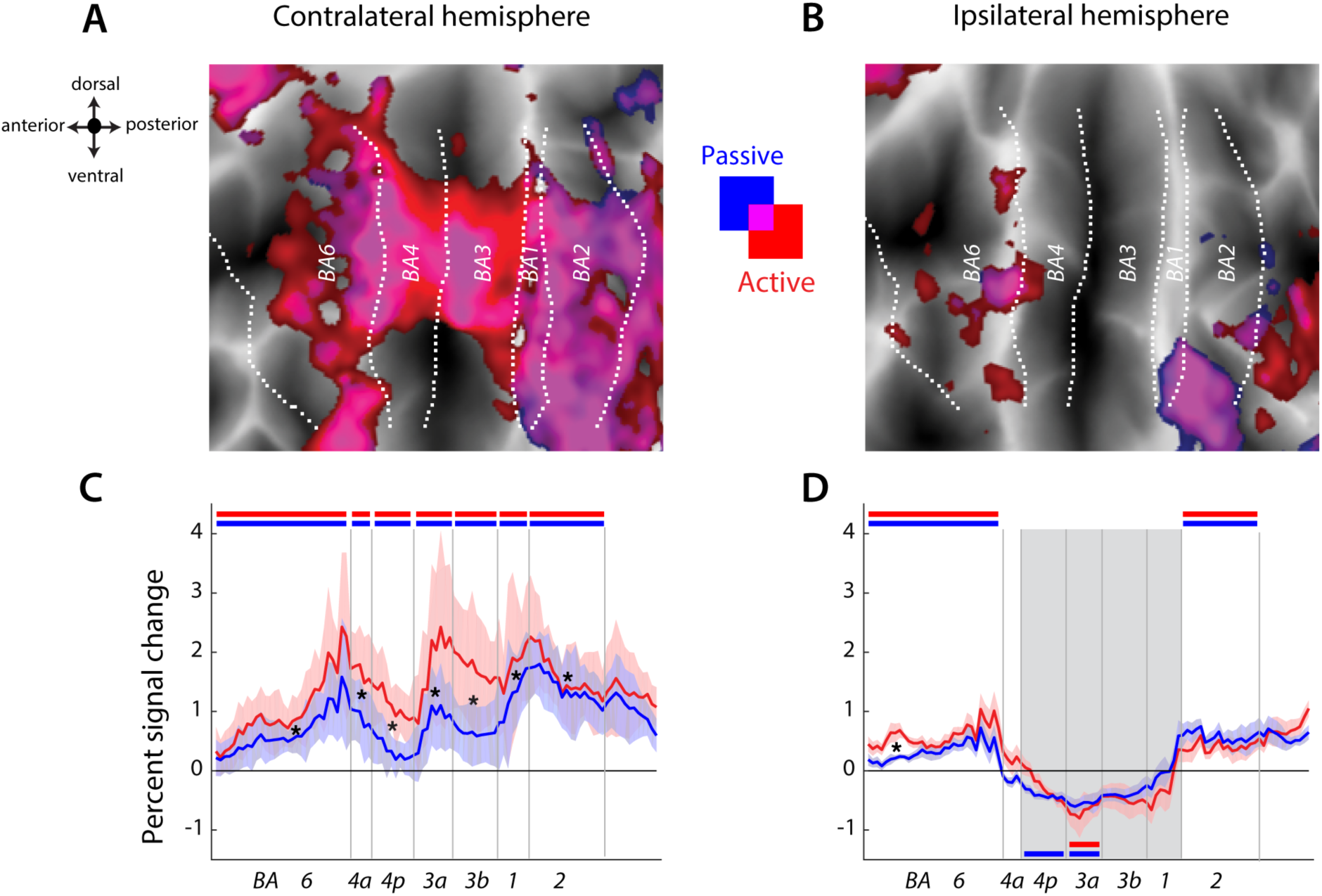
Average evoked activation during active and passive tasks across subfields of sensorimotor cortex of the contra- and ipsilateral hemisphere. (**A**) Evoked activity for the active (red) and passive (blue) conditions on the flattened contralateral cortical surface. The two conditions activated similar cortical areas, with the overlap indicated by purple areas. Regions of interest (ROIs) were defined based on the probabilistic cyto-architectonic atlas (Fischl et al., 2008), with each node assigned the area of the highest probability. Borders between regions are indicated with white dotted lines. (**B**) Evoked activity above resting baseline for the two conditions on the flattened ipsilateral hemisphere. (**C**) Percent signal change for active and passive tasks was sampled in a cross-section from anterior (BA6) to posterior (BA2), along a rectangular strip with a width of 26 mm. Horizontal red and blue bars indicate significant activation during the active and passive task, respectively. Significant differences between the activation for active and passive tasks are indicated by black stars (p<0.0071 – Bonferroni correction). (**D**) Ipsilateral hemisphere showed suppression of activity below resting baseline around the central sulcus for both conditions, indicated with grey background. BA6 displayed more activation for the active than passive condition, but all other areas responded similarly for the two conditions.

Next, we evaluated how strong representations for different fingers were in each of these ROIs, independent of the overall activity. It is possible to observe large activation without any representation of individual fingers (implying the activation is induced by processes not specifically related to finger control), or to observe lower activation with very clear finger representation. For a region to perform a specific function, a clear representation is more important than high activation (Diedrichsen and Kriegeskorte, 2017). We evaluated the strength of representation using the cross-validated squared Mahalanobis distance estimate (crossnobis, Diedrichsen et al., 2016) between activity patterns of individual fingers, separately for active presses and passive finger stimulation. As expected, we found strong finger representations for both passive and active conditions (Fig. 3A), confirmed by a t-test on distance estimates of each condition across all cortical sensorimotor regions combined (passive: *t*_(6)_=13.82, *p*=8.93e^-6^, active: *t*_(6)_=9.76, *p*=6.65e^-5^). Distances were particularly large in the depths of the central sulcus, peaking in area 3b, and decreased anteriorly in premotor area (BA6) and posteriorly in BA2 (Fig. 3C). We quantified this observation statistically by performing a condition x ROI ANOVA on the distance estimates. The main effect of condition was not significant (*F*_(1,6)_=3.183, *p*=0.125), but both the main effect ROI and the interaction between the ROI and condition were (ROI: *F*_(1,6)_=37.288, *p*=5.1e^-14^, interaction: *F*_(6,36)_=12.183, *p*=1.9e^-7^). Post-hoc t-tests on the effect of condition within each region revealed a trend for larger distances in the passive compared to active condition in BA3b and BA1, but this difference did not reach significance after Bonferroni correction.

**Fig. 3.**
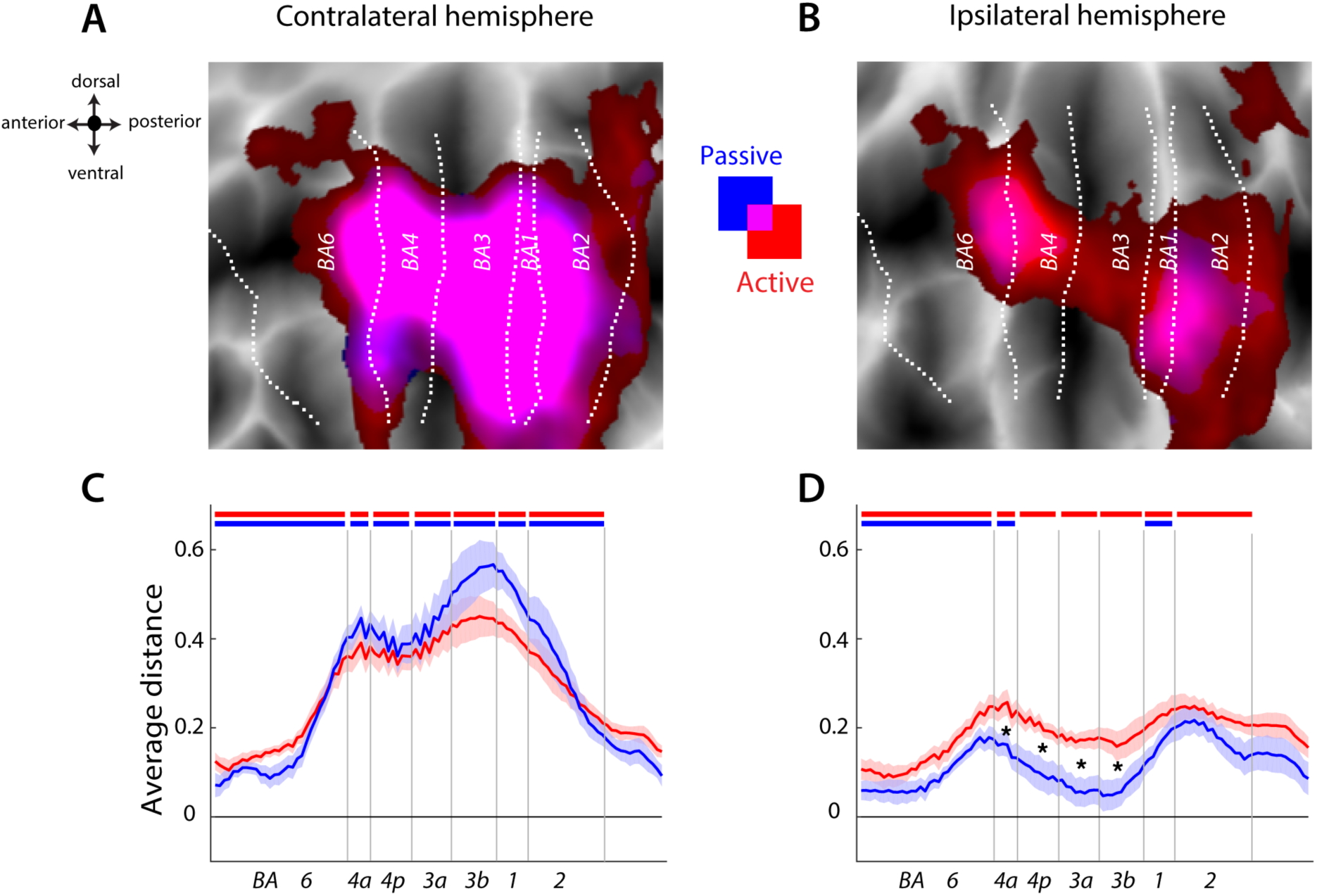
Average distance between finger patterns for active and passive conditions across the two hemispheres. (**A**) Average distance between finger patterns for the active (red) and passive (blue) tasks on the flattened contralateral cortical surface. The two conditions evoked similar distances, which is indicated by the purple overlap. (**B**) Average passive and active distances in the ipsilateral hemisphere. The active condition elicited higher distances than the passive condition, which is reflected in the predominately red blobs, especially in the depth of the central sulcus. (**C**) Distances in the contralateral hemisphere were significantly higher than zero for active and passive tasks, as indicated by the red and blue bars. There was no difference in distances between the two conditions in any region of interest (ROIs). (**D**) Ipsilateral hemisphere displayed higher distances for the active than the passive task. This difference was significant in areas BA4a, 4p, 3a and 3b (asterisks, p<0.0071).

In summary, we found that both active and passive conditions activated the finger-specific representations to the same extent in contralateral M1 and S1. In contrast, the average overall activity was significantly higher in the active condition. This means that the additional neuronal processes in the active condition were not finger specific, but instead increased activity in a general fashion for all fingers.

### Active and passive condition elicit largely shared patterns in the contralateral hemisphere

The strength of the representation of individual fingers in the contralateral hemisphere was comparable for active finger presses and passive finger stimulation. But do active and passive conditions activate the same circuits? On one extreme, individual finger movements and individual finger stimulation could evoke the same responses in the same voxels. On the other extreme, the two conditions could activate completely different voxels or the same voxels to different extents. Using PCM, we can determine the degree to which finger-specific patterns of activity were shared across the two conditions. When estimating the correlation between active and passive conditions (corrected for the measurement noise, see methods), we obtained an average value of 0.84 between active movement and sensory stimulation across all areas of interest (Fig. 4A). The correlation coefficient was the highest in the BA1 with *r*=0.91 (one-sample t-test of subject-specific estimates against zero: *t*_(6)_=33.60, *p*=2.3e^-8^), next in the neighbouring primary motor cortex (*r*=0.87, *t*_(6)_=21.72, *p*=4.7e^-8^) and BA3 (*r*=0.86, *t*_(6)_=21.72, *p*=3.1e^-7^). The correlation was above 0.8 also in the premotor cortices (*r*=0.81, *t*_(6)_=18.46, *p*=8.1e^-7^). BA2 displayed the lowest correlation (*r*=0.75, *t*_(6)_=19.33, *p*=6.2e^-7^) amongst chosen ROIs.

**Fig. 4.**
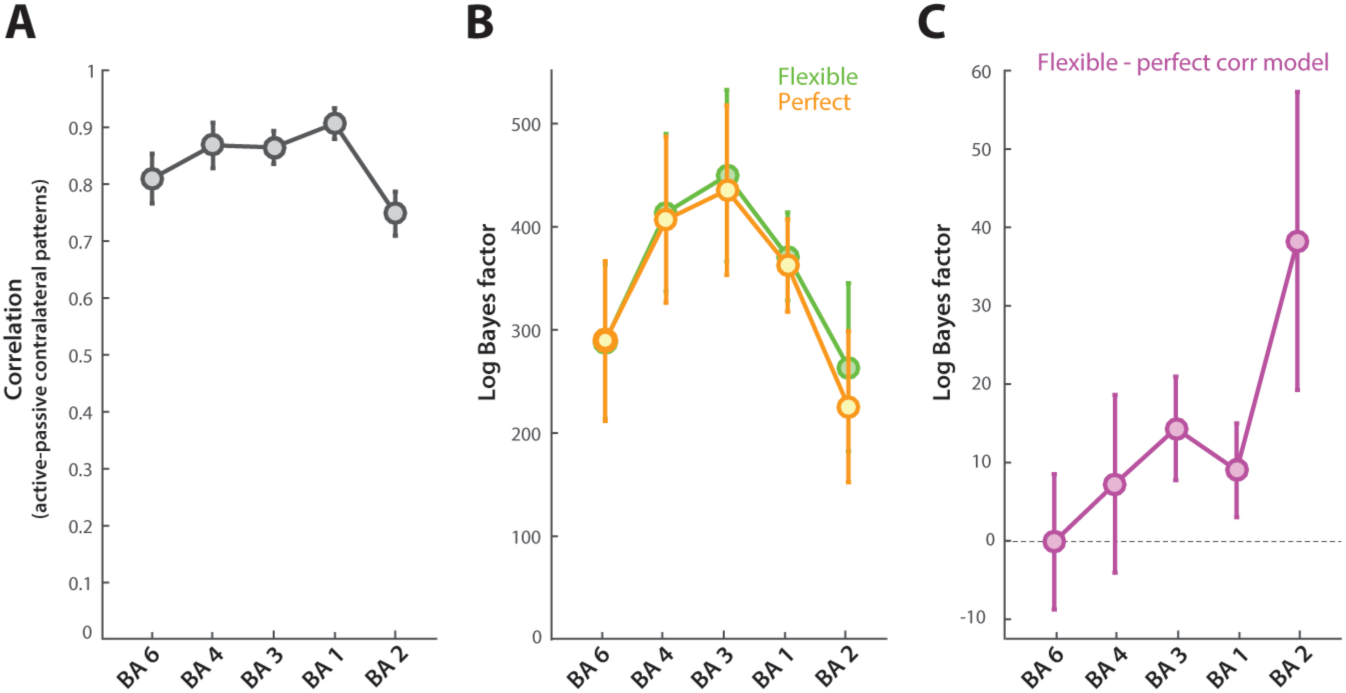
Correlation between contralateral activity patterns evoked by single finger presses and sensory stimulation. (**A**) Correlations were estimated using pattern-component modelling (PCM). (**B**) Performance of the model with correlation between active and passive patterns unconstrained (‘flexible correlation’ model) and the model where the correlation is constrained to be one (‘perfect correlation’ model) – both expressed relative to a ‘null’ model (no correlation between active and passive patterns). (**C**) Direct comparison of ‘flexible’ correlation and ‘perfect correlation’ models, assessed by the difference in their log Bayes factors. While there was on average evidence for the flexible model to be better, this difference was not significant across subjects.

These results clearly show that the passive and active conditions engage highly overlapping finger-specific circuits. Given that the correlation estimates are smaller than 1, there may be some specific difference in the patterns evoked by sensory and motor conditions. However, the problem is that correlation coefficients underestimate the true correlation (Diedrichsen et al., 2017). To test whether the data could be explained by a true correlation of *r*=1 between active and passive patterns, we compared two PCM models: a *‘perfect correlation’ model* which constrained the correlation between passive and active patterns to 1, and a *‘flexible correlation’ model* in which the correlation was estimated in a cross-validated fashion across subjects. Evidence for these two models was expressed relative to a *‘null’ model* which assumes that the correlation between active and passive patterns is 0. Both flexible and perfect correlation models were a better descriptor of our data than the null model (Fig. 4B) – the *flexible correlation model* had a log-Bayes factor of 357 (one-sample t-test against zero: *t*_(6)_=10.684, *p*=3.41e^-16^), whereas the *perfect correlation model* had a log-Bayes factor of 344 (*t*_(6)_=10.188, *p*=2.53e^-16^). We next compared the performance of the two models across ROIs (Fig. 4C), using an ROI x model ANOVA. Neither the main effect of the region nor the model type were significant (ROI: *F*_(4,24)_=2.567, *p*=0.064; model: *F*_(1,6)_=1.941, *p*=0.213). The interaction between the two factors was significant (*F*_(4,24)_=4.773, *p*=0.0056), but post-hoc t-tests revealed no significant difference in model performance in any of the regions. This suggests that a model that assumes that passive and active patterns are perfectly correlated explains our data as well as one that freely estimates the correlation. Altogether, our results demonstrate that individual finger movement and sensory processing activate virtually the same finger-specific patches of contralateral sensorimotor cortex, without consistent evidence that would suggest separate activation patterns.

### Ipsilateral finger representations are stronger in active than passive condition

Having quantified the similarity of passive and active digit representations in the contralateral hemisphere, we next turned to the ipsilateral hemisphere. We again first quantified the overall percent signal change of elicited activity. Consistent with previous research (Verstynen et al., 2005; Diedrichsen et al., 2013) we found significant BOLD modulation across ipsilateral ROIs during the active condition, as confirmed by a one-way ANOVA with the main effect of region (*F*_(6,36)_=16.26, *p*=5.9e^-9^). Activation in the depth of the sulcus was suppressed below resting baseline (Fig. 2D, grey background), and this suppressive effect was significant in areas 4p and 3a (BA4p: *t*_(6)_=-4.89, *p*=0.0027; BA3a: *t*_(6)_=-4.28, *p*=0.005). Only premotor (BA6) and parietal areas (BA2) exhibited significant increases in BOLD signal (BA6: *t*_(6)_=6.57, *p*=5.94e^-4^; BA2: *t*_(6)_=4.51, *p*=0.004).

To quantify the activation and deactivation profiles across both active and passive conditions, we used a condition x ROI ANOVA. The main effect of condition was not significant (*F*_(1,6)_=0.095, *p*=0.769), but there was a significant main effect of ROI (*F*_(1,6)_=19.55, *p*=5.4e^-10^), and a significant interaction between the two factors (*F*_(6,36)_=8.13, *p*=1.4e^-5^). Post-hoc t-tests demonstrated that this interaction was driven by higher activity in the premotor cortex during the active condition (*t*_(6)_=4.23, *p*=0.006), which is in line with its bilateral involvement during action preparation (Cisek et al., 2003). Other areas showed no significant difference in activity between the two conditions. Thus, regions located in the depth of the central sulcus in the ipsilateral hemisphere were significantly suppressed during both passive and active conditions.

We have previously found that despite the suppression of BOLD activity, the ipsilateral hemisphere contains information about individual finger movements (Diedrichsen et al., 2013). Here we asked whether the ipsilateral hemisphere represents individual fingers only during active movement, or also during passive finger stimulation. We first examined individual finger representations during the active condition. The average distance among active finger presses was higher than zero in every region (all *t*_(6)_>2.721, *p*<0.034; Fig. 2D), replicating our prior results (Diedrichsen et al., 2013). Next we tested whether the ipsilateral hemisphere represents individual fingers in the passive condition to the same extent as during active movement (similar to the contralateral hemisphere). The main effect of condition on the distance estimates was significant (*F*_(1,6)_=24.36, *p*=0.0026), and post-hoc t-tests revealed that the average distance was lower in the passive than the active task in the depth of the sulcus (black stars in Fig. 2D). Subfields 4p, 3a, and 3b all showed significant distances during active finger presses, but did not show finger representation for passive finger stimulation, as confirmed by one-sample t-tests against 0 (no blue bars in Fig. 2D). These findings suggest that ipsilateral representations in these areas are driven by processes involved in the active generation of movement, but not by the sensory input arising from passive movement.

Last, we quantified whether the relative amount of finger representation across the passive and active tasks differs across the two hemispheres. This test is critical to determine whether the source of contralateral and ipsilateral finger information is identical or different. A hemisphere x condition ANOVA across all regions combined revealed a clear interaction effect (*F*_(1,6)_=64.481, *p*=2.0e^-4^), demonstrating that the relative magnitude of finger-specific representation during the active and passive conditions differs significantly across the two hemispheres. While the contralateral sensorimotor circuit represents individual finger presses and stimulation to the same extent, finger representation on the ipsilateral side was stronger during the active condition. This demonstrates that the contribution of sensory information to the neural activation patterns is much smaller in the ipsilateral as compared to the contralateral sensorimotor areas.

### Spatial distribution of active representations is different across hemispheres

An additional important insight about ipsilateral representation can also be gained by considering the spatial distribution of representations across subfields of sensorimotor cortices. We compared the distribution of active distances across the cross-section of ROIs in the contralateral and ipsilateral hemisphere (i.e. the profile of red lines in Fig. 2C versus Fig. 2D). Our results showed that the ipsilateral profile of distances for the active condition is not just a scaled-down version of the contralateral distances. For example, contralateral distances peaked in area 3b, but ipsilateral hemisphere showed lower distances in 3b than in areas 1 and 2. To quantify this effect, we performed a hemisphere x ROI ANOVA on the distance estimates in the active condition. Both main effects were significant (hemisphere: *F*_(1,6)_=35.827, *p*=0.001, ROI: *F*_(6,36)_=20.272, *p*=3.33e^-10^), but importantly the interaction between them was significant as well (*F*_(6,36)_=17.236, *p*=2.83e^-9^). This suggests that the ipsilateral hemisphere has a unique profile across areas, with relatively stronger finger-specific representations in premotor and parietal areas.

## Discussion

In this study, we used active finger presses and passive finger stimulation to investigate the origin of finger representations in ipsilateral sensorimotor cortex. We first provided a detailed characterization of the nature of contralateral representations during active and passive conditions. We found that despite BOLD-activations being larger for the active condition, the corresponding finger-specific representations were equally strong across the two conditions. We expanded upon these results in two ways. First we demonstrate that finger-specific activity patterns were highly correlated between active and passive conditions. Indeed, considering the measurement noise, there was no evidence that passive and active patterns differed in any way.

Second, we quantified finger representations across the subfields of the sensorimotor cortex, and report that representations were most pronounced in BA3a, 3b and BA1. Altogether, our results demonstrate that passive finger stimulation drives contralateral finger-specific motor circuits as strongly as active finger presses, emphasizing the importance of sensory inputs in dexterous feedback control (Pruszynski et al., 2016).

Having established the nature of contralateral sensorimotor finger representations, we then examined the extent to which the ipsilateral motor areas were recruited during active and passive conditions. Overall, ipsilateral representations were weaker than those in the contralateral hemisphere. Critically, however, while contralateral representations were equally strong for both active and passive conditions, ipsilateral representations were significantly stronger for the active condition. Moreover, there was no reliable finger representation during passive stimulation in ipsilateral areas 4p, 3a and 3b. The difference between hemispheres became also clear when investigating the spatial distribution of finger representations across the two hemispheres – on the contralateral side, finger representations were strongest along the central sulcus, whereas on the ipsilateral side, they were strongest in premotor and parietal areas. Together, these findings suggest that ipsilateral sensorimotor representations qualitatively differ from their contralateral counterparts. This implies that ipsilateral representations are not a pure consequence of trans-callosal spill-over from the homologous areas (e.g. M1-M1; Asanuma and Osamu, 1962), otherwise ipsilateral representations should have been equally strong in the active and passive conditions.

One possible explanation for these differences could be that ipsilateral representations are more related to the planning and initiation of actions, but less to the ongoing feedback control of movements. This proposed role of ipsilateral representations in movement planning is consistent with research in non-human primates demonstrating that the ipsilateral hemisphere has limited access and capacity to causing upper-limb movements (Kuypers et al., 1962). In particular, stimulation of the ipsilateral pathway does not produce monosynaptic and/or disynaptic activity in the ventral horn neurons (Soteropoulos et al., 2011), suggesting that these ipsilateral pathways play at most a modulatory role in feedback control. Previous studies have highlighted the fact that the ipsilateral sensorimotor areas have a strong representation of the position, velocity, or direction of upper limb movement (electrophysiology: Donchin et al., 2001; Ganguly et al., 2009; fMRI: Verstynen et al., 2005; Diedrichsen et al., 2013; Haar et al., 2017; ECoG: Scherer et al., 2009; Fujiwara et al., 2017). Our results now suggest that even if qualitatively similar representation is observed on both sides, the underlying neurophysiological origins may differ, with the ipsilateral representations stemming from bilateral planning processes, while the contralateral hemisphere are directly related to feedback control.

Importantly, however, we cannot discern whether finger representations in ipsilateral primary sensorimotor areas have functional relevance, or whether they are a pure epiphenomenon. Namely, the presence of a detailed representation during the active condition (as observed with fMRI or electrophysiology) does not automatically imply that the activity plays any causal role in the planning of the movement. For example, bilateral representations in primary sensorimotor regions could arise from covert planning of candidate responses with either hand (Cisek and Kalaska, 2010). The ipsilateral representations would then be suppressed when the choice of hand is made, without contributing in any way to motor performance. While there is some evidence that disruption of ipsilateral motor circuits impede the quality and skill of motor execution (Chen et al., 1997), the observed deficits are rather subtle (Noskin et al., 2008; Xu et al., 2017).

An important insight into the nature of ipsilateral representations comes from an optogenetic experiment in mice, which demonstrated that disruption of ipsilateral premotor cortex did not lead to direct behavioral deficits (Li et al., 2015). However, the ipsilateral motor areas were able to compensate for silencing of corresponding motor areas in the contralateral hemisphere. Thus, ipsilateral representations may not be essential during normal function, but may assume a compensatory function in the case of disruption (Johansen-Berg et al., 2002).

In conclusion, we have provided a detailed characterization of the nature of ipsilateral sensorimotor representations during active presses and passive finger stimulation. Our results suggest that the ipsilateral hemisphere does not receive the sensory input critical for dexterous feedback control, and instead may primarily be involved in planning-related processes. Therefore, our study provides important constrains on the role that the ipsilateral hemisphere can play in the control of movement in health and disease.

## Acknowledgements

This work was supported by a James S. McDonnell Foundation Scholar award, an NSERC Discovery Grant (RGPIN-2016-04890), and the Canada First Research Excellence Fund (BrainsCAN). The authors wish to thank Tamar Makin for comments on the early version of the manuscript.

